# Systematic analysis of cell phenotypes and cellular social networks in tissues using the histology topography cytometry analysis toolbox (histoCAT)

**DOI:** 10.1101/109207

**Authors:** Denis Schapiro, Hartland Warren Jackson, Swetha Raghuraman, Vito R T Zanotelli, Jana R Fischer, Daniel Schulz, Charlotte Giesen, Raúl Catena, Zsuzsanna Varga, Bernd Bodenmiller

## Abstract

Single-cell, spatially resolved ‘omics analysis of tissues is poised to transform biomedical research and clinical practice. We have developed an open-source computational histology topography analysis toolbox (histoCAT) to enable the interactive, quantitative, and comprehensive exploration of phenotypes of individual cells, cell-to-cell interactions, microenvironments, and morphological structures within intact tissues. histoCAT will be useful in all areas of tissue-based research. We highlight the unique abilities of histoCAT by analysis of highly multiplexed mass cytometry images of human breast cancer tissues.

Technological advances in the multi-parametric analysis of single cells have revealed an unprecedented heterogeneity of cellular phenotypes and functional states that are concealed in population-based studies^1–3^. Each cellular phenotype is defined by the interplay of its internal state and the environment in which it resides, and tissue and organ function is the output of these coordinated cell activities. Deregulation of inter-cellular communication is central to many diseases such as cancer. Consequently, the ability to analyze functional states on the single-cell level with spatial resolution is key to understanding normal tissue function, disease biology, and for the development of treatments of disease^4–6^.

Recent techniques such as FISSEQ^7^, MERFISH^8^, cycling immunofluorescence^9–11^, multiplexed ion beam imaging (MIBI)^12^, and imaging mass cytometry (IMC)^13^ allow for single-cell, spatially resolved, highly multiplexed analysis of solid tissues and provide essential information including the distribution of transcripts, proteins, and their modifications within single cells, microenvironments, and entire tissues. Despite these experimental advances no computational approach has been developed to enable the comprehensive and quantitative interactive exploration of spatially resolved, highly multiplexed tissue measurements. Current open-source tools that provide image-linked data analysis are typically focused on the analysis of cell lines imaged with low-plex fluorescence microscopy or basic tissue histology and are not geared to the analysis of highly multiplexed measurements^14–16^. On the other hand, tools that have been developed to perform analyses of non-imaging highly-multiplexed single cell data (e.g., suspension based mass cytometry) do not exploit spatial information (Supplementary Fig. 1)^17,18^.

In order to provide a complete picture of a tissue ecosystem, define molecular and spatial signatures necessary for analysis of tissue biology, and, in the case of disease, identify clinically relevant features, it is necessary to analyze and interrelate layers of information obtained from molecular measurements on cells, cell populations, cell-to-cell interactions, microenvironments, tissues, and experimental cohorts. Here we present a powerful, interactive computational platform that we call histoCAT that makes quantitative analysis of highly multiplexed, single-cell-resolved measurements possible. histoCAT combines intuitive high-dimensional image visualization, state-of-the-art analysis methods for cell phenotype characterization, and novel algorithms for the comprehensive study of cell-to-cell interactions and the “social” networks of cells within complex tissues (Fig. 1). This provides unprecedented capabilities for investigators from biology, biomedicine, and pathology to investigate tissue changes during health, disease, and treatment.

**Figure 1.**
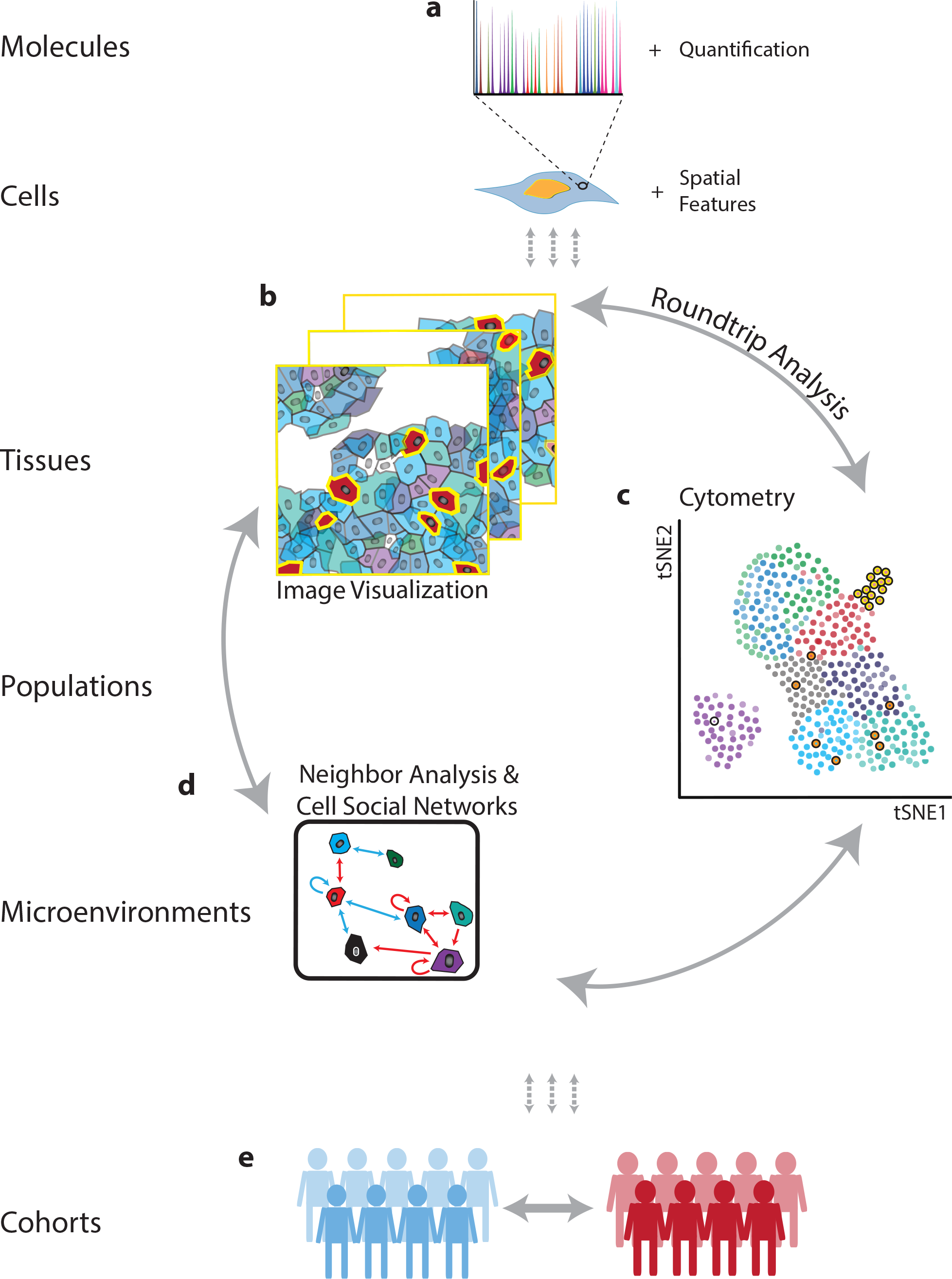
From molecular to clinical information: multi-scale analysis of the tissue ecosystem.(**a**) Spatially resolved, high-dimension molecular measurements are aggregated using image masks to define regions corresponding to each cell. (**b**) Visualization of images, (**c**) cytometry analysis and (**d**) analysis of neighbors and cellular interaction networks facilitate “round-trip” analysis through layers of information. (**e**) Using molecular, cellular and spatial signatures, experimental cohorts can be compared and contrasted.

In histoCAT, all single cell information, including spatial features (Fig. 1a), is linked to the corresponding multiplexed image enabling visualization of images and single-cell analysis in parallel (Fig. 1b, c). histoCAT uses images and a corresponding segmentation mask to extract single-cell data including abundances of all measured markers for a cell and area of interest, as well as spatial features (like cell size and shape), and aspects of the cell’s environment such as cell neighbors and cell crowding. This information is compiled into a flow cytometry standard format (.fcs) file for further analysis inside or outside of histoCAT. “Round-trip” analyses from a specific area of an image to dataset-wide analyses of single-cell phenotypes and their interactions and back to the visualization of unique cells on the images enables users to define and understand key cell populations and their spatial context in the tissue. To enable the quantitative and systematic analysis of the “social” networks of cells, we developed a novel algorithm to identify all direct cell-cell neighboring interactions (Fig. 1d) and determine the significant interactions and unique cell environments across entire datasets and within specific cohorts (Fig. 1d, e).

To combine image-based spatial information and high-dimensional cytometry data, the histoCAT GUI is divided in two parallel sections for paired image and cytometry analysis (Supplementary File). In the image visualization section of histoCAT, high-dimensional images (Fig. 2a) as well as cell masks, single-cell marker quantification, and cell identification labels can be visualized. In the analysis section of histoCAT, image-derived marker quantification and spatial features of single-cell data are extracted for each image (Fig. 2a), combined (Fig. 2b), and visualized using multi-dimensional reduction tools such as t-SNE^17^ maps (Fig. 2b), scatter plots, histograms, box plots, or other visualizations (Supplementary File).

To demonstrate the potential of histoCAT-powered analyses, we investigated the cellular phenotypes and cellular microenvironments of human breast cancer as visualized by IMC. By pairing classic immunohistochemistry staining, high-resolution tissue laser ablation, and mass cytometry, IMC can measure abundances of more than forty unique metal-isotope-labeled tissue bound antibodies simultaneously at a resolution comparable to fluorescence microscopy in a single tissue section^13^. Fifty-two diverse breast cancer samples were stained with an antibody panel designed to enable identification of cell lineages and detection of signaling pathway activation, proliferation, apoptosis, and clinical markers (Supplementary Table 1 and Supplementary Table 2).

To gain a tissue-wide overview of the cell phenotypes present in a given image set, we have incorporated two approaches into histoCAT. The first approach is supervised and based on tSNE^17^, a data dimensionality reduction approach that projects cell phenotypes defined by a multitude of markers into two dimensions, grouping highly similar cells (Fig. 2a-c). Expression of individual markers can be highlighted using color scales, manually gated, and annotated for the corresponding cell phenotype (Fig. 2d). The second approach is based on the unsupervised clustering algorithm PhenoGraph^19,20^. PhenoGraph identified 29 phenotypes shared across images and clinical subgroups, which were then visualized on a tSNE map (Fig. 2c, Supplementary Fig. 2). These phenotypes were characterized by specific epitopes (e.g., vimentin, phenotype #4, and CD68, phenotype #7) and combinations of markers (e.g., proliferative Ki-67-positive and phospho-S6-positive phenotypes #8, #10, and #19) (Fig. 2e). Any of these cell populations can be linked back to their source images to visualize cells within the context of the multi-cellular environment (Fig. 2f).

**Figure 2.**
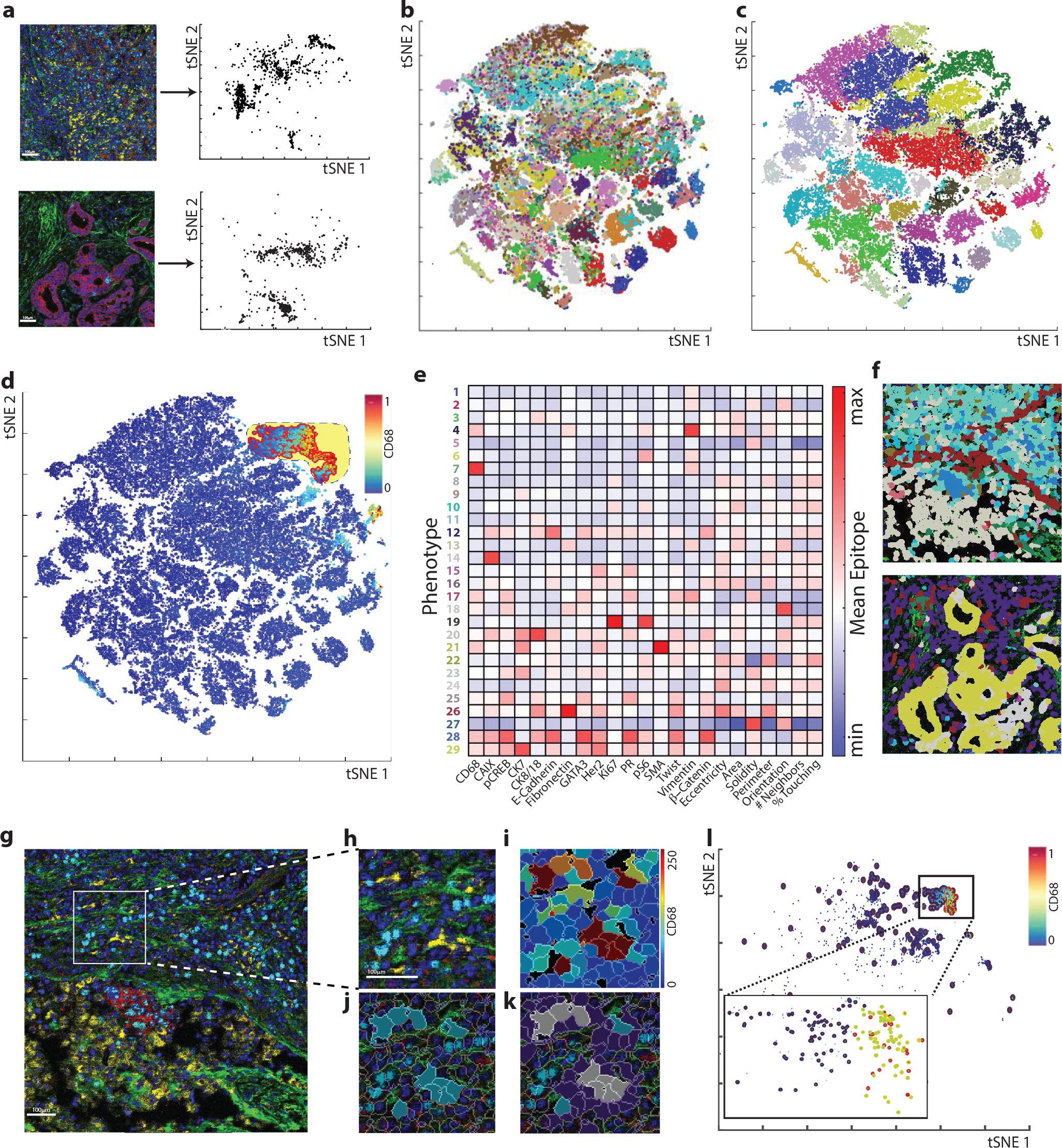
Round-trip analysis of unique cell types in high-dimension images of breast cancer. (**a**) Multi-parametric images are displayed in the histoCAT image window using up to six user-defined color channels (red, E-cadherin; green, vimentin; blue, histone H3; cyan, Ki-67; magenta, cytokeratin 7; yellow, CD68). Scale bar = 100 µm. High-dimension single-cell data, including spatial features and all expressed markers from each segmented cell are extracted from each individual image and visualized in a t-SNE plot. (**b**) When combined, distinct colors distinguish cells of each source image when an entire dataset is visualized in one t-SNE plot. (**c**) Unsupervised clustering of all cells according to their marker expression throughout the dataset using PhenoGraph defines complex cell phenotypes and enables labeling of each cell phenotype cluster with a distinct color. (**d**) Alternatively, quantification of an individual parameter can be heatmapped onto the t-SNE plot and populations can be identified in a supervised manner using the gating tool. (**e**) Cell phenotypes can be further investigated using plotting tools such as heatmaps. (**f**) Single cells are colored according to the identified phenotypes within the context of the tissue microenvironment on their original image. (**g**) All images containing cells of a subpopulation of interest can then be identified and loaded: red, E-cadherin; green, fibronectin; blue, histone H3; cyan, Ki67; magenta, cytokeratin 7; yellow, CD68. Scale bar = 100 µm. Images can be visualized using (**h**) pseudo-color or by (**i**) heatmap representing the intensity of a marker in each cell. (**j**) Cells of interest can be highlighted on the image (turquoise) and neighboring cells (purple or gray if representing both subpopulations) within a defined pixel range can also be identified and highlighted on (**k**) the image or (**l**) the analysis plot of the individual image (red highlight, cell of interest; blue highlight, neighbor; yellow highlight, both subpopulations).

Tumor associated macrophages (TAMs) are a key cell type of the tumor microenvironment. Depending on their polarization state, TAMs can drive or hinder tumor progression and thus are highly attractive biomarkers and drug targets^21^. To gain a deeper understanding of TAMs and their neighborhoods, we used the expression of CD68 to select macrophages by gating on the tSNE plot (Fig. 2d, Supplementary File) or alternatively selecting the CD68^+^ PhenoGraph phenotype #7 (Fig. 2e). Macrophage CD68 epitope expression can be visualized in images by color (yellow, Fig. 2g,h), heatmapped for each segmented cell (Fig. 2i), or CD68^+^ cells selected from a plot can be highlighted on source images in their original tissue context (Fig. 2j).

To investigate the microenvironment histoCAT has two neighborhood functions. The first function is user guided and returns a subpopulation of cells touching or proximal to a cell population of interest for visualization on images (Fig. 2k) or in downstream analysis (Fig. 2l). This analysis showed that distinct proliferative (Ki-67, phospho-S6) and hypoxic (carbonic anhydrase IX) epithelial tumor cells neighbor CD68^+^ macrophages (Supplementary Fig. 3). The second neighborhood function enables the unbiased and systematic study of all cell-to-cell interaction patterns present in a tissue or all tissues of a sample cohort. Such unbiased analyses are needed to generate novel biological hypotheses or to identify novel clinically relevant features. histoCAT enables such a comprehensive neighborhood analysis using a permutation test to compare the number of interactions between cell types in a given image to that of a matched control containing randomized/shuffled cell phenotypes (Fig. 3a). This approach determines the significance of cell-to-cell interactions and reveals enrichments or depletions in cell-to-cell interactions that are indicative of cellular organization within a tumor. As an output of the comprehensive neighborhood analysis, the significance of a neighboring interaction between every pair of phenotypes is visualized as a heatmap in which each row represents the neighborhood of a cell phenotype of interest and every column the enrichment or depletion of another cell in its neighborhood. This type of analysis was performed for data obtained on the 52 breast cancer samples (Fig. 3b). These directional interactions identify cells surrounding or being surrounded by another cell type. For example, the highly interactive tumor cell phenotype #3 is surrounded by stromal cell phenotype #13, but phenotype #3 is not significantly enriched in the surroundings of phenotype #13 (Fig. 3b, red squares and blue squares respectively, Supplementary Fig. 4). For TAMs (phenotype #7) this unsupervised neighbor analysis revealed significant interactions with the tumor cell phenotype #22 (Fig. 2e; Fig. 3b, row 7 column 22) identifying a key subset of TAM interacting cells within all neighbor interactions (Supplementary Fig. 3j). This suggests a relationship and cellular crosstalk between CD68^+^ cells and E-cadherin^+^/phospho-S6^+^/ Twist^+^ tumor cells and identifies a distinct tumor microenvironment for future study.

**Figure 3.**
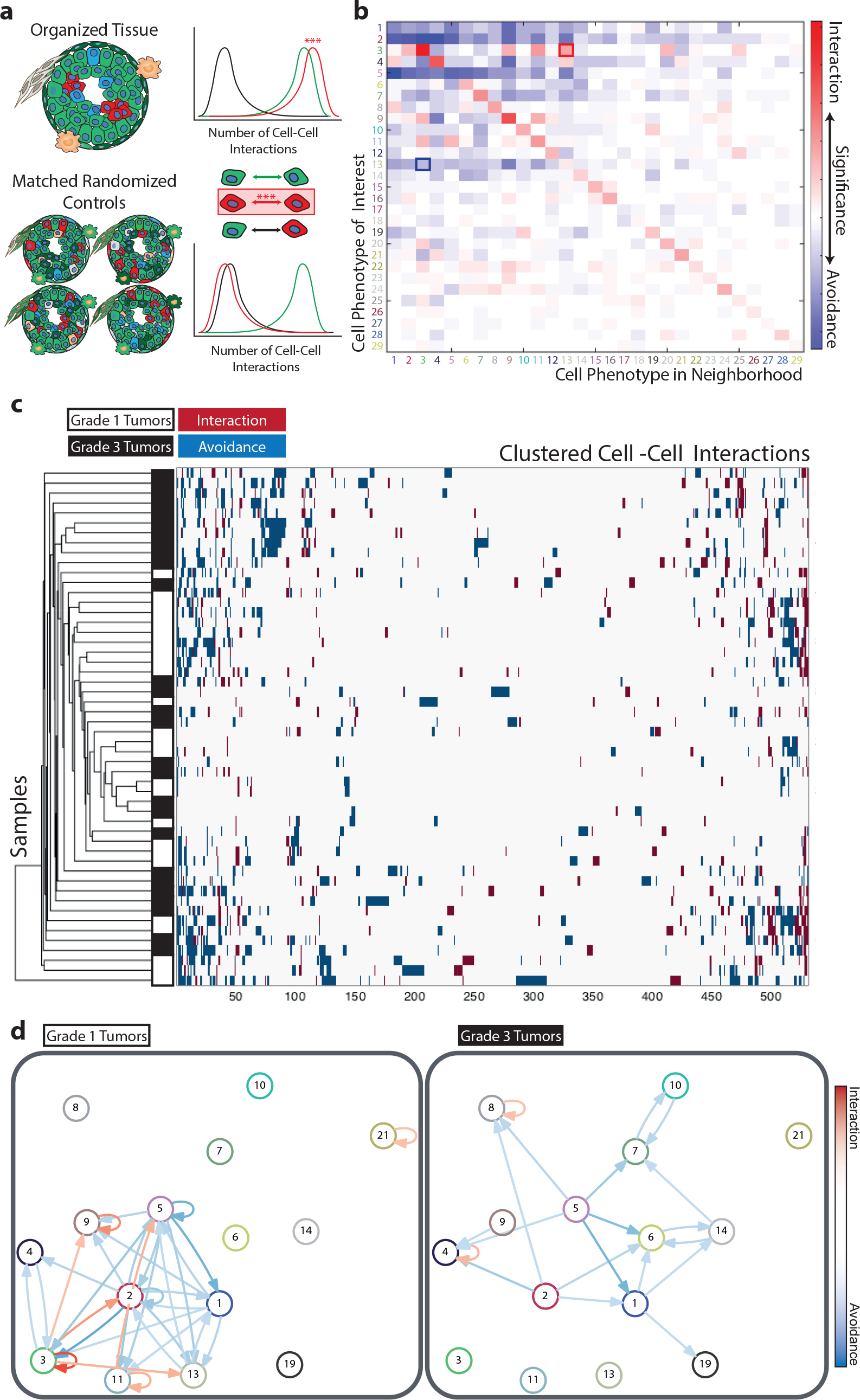
Neighborhood analysis of breast cancer cell phenotypes. (**a**) Schematic of neighbor analysis in which the prevalence of a particular cell-cell interaction in an image is quantified and significance is determined by comparison to its prevalence in cell-type-randomized controls of the same image. Number of interactions between abundant cells (green), between rare clustered cells (red), and between abundant cells and rare cells (black). (**b**) Every cell interaction present in more than 10% of all images plotted as a heatmap in which square color indicates whether the cell type in the row is significantly neighbored (red) or avoided (blue) by the cell type in the column (p < 0.01). Highlighted squares indicate an example of a directional interaction where stromal phenotype #13 surrounds tumor cell type #3 (red square), but #3 is not surrounded by #13 (blue square). (**c**) Clustering of all samples and cell-cell interactions according to the presence of significant (p < 0.01) phenotype interaction (red) or avoidance (blue). White represents interactions that are not present or not significant. (**d**) Force-directed cell interaction network graphs representing the organization of PhenoGraph-defined cell phenotypes in Grade 1 and Grade 3 tumors. Circle color corresponds to PhenoGraph cluster, arrow color indicates the direction of the interaction (red interaction, blue avoidance), and intensity of the line color indicates significance. A connection is only visualized if the interaction or avoidance is significant in a least 20% of the grouped samples and the cell phenotypes were simultaneously present in at least 40% of the grouped samples.

In a next step, we investigated whether we could identify patterns of cell-to-cell interactions that relate to the available clinical data of our patient cohort. We clustered all images based on their significant cell-to-cell interactions (Fig. 3c), and a clear grouping of Grade 1 and Grade 3 tumors became apparent. This separation was driven by dense interaction of tumor cell phenotypes in Grade 1 tumors, and segregated hypoxic (phenotype #14) and proliferative cell phenotypes (phenotype #10 or #19) in more advanced Grade 3 breast cancers (Fig. 3c, Supplementary Fig. 4, Supplementary Fig. 5a). Thus, clustering of images based on significant cell interactions defines groups of tissues (in this case cancer samples) that have similar organization (Fig. 3c).

To facilitate the visualization and comparison of cell-to-cell interactions over large datasets, we used our neighbor algorithm to detect cell social networks. The resulting cell interaction/avoidance networks of Grade 1 or Grade 3 samples revealed tumor-grade specific multi-cell interaction/avoidance clusters (Fig. 3d). Different clusters of tumor cells (phenotype #3, #9, or #1) were neighbored by tumor cells of other phenotypes and/or stromal cells (phenotype #2, #5, #13) within Grade 1 but not Grade 3 samples (Fig. 3d, Supplementary Fig. 5). Overall, fewer significant interactions were observed in the more advanced and less organized Grade 3 tumors than in Grade 1 tumors (Fig. 3d, Supplementary Fig. 5). Thus, cellular organization and distinct ecosystems are present in specific clinical settings and could be used to distinguish unique disease states.

By combining cytometry and image analysis and novel algorithms for cell-to-cell interaction network analysis within one toolbox, histoCAT is able to define complex cell types using multiplexed measurements and spatial features as parameters and to elucidate patterns of cellular interactions within heterogeneous tissues. The use of round-trip analyses between single-cell data and source images using machine learning and community-finding algorithms within an intuitive user interface will enhance our understanding of tissue structure at the cellular level. Combined with focused, hypothesis-driven datasets, future investigations of multiplexed imaging cytometry data using histoCAT could reveal cell types and cell interactions that drive disease. histoCAT is open source, and we invite the community to further develop this tool for the analysis of next-generation imaging and pathology data.

## Data availability

All raw data and software can be downloaded upon publication of the manuscript at https://github.com/BodenmillerGroup/histoCAT and http://www.bodenmillerlab.org/research-2/histoCAT/

## ACKNOWLEDGEMENTS

We would like to thank the Bodenmiller lab for support and fruitful discussions. This work was supported by the Swiss National Science Foundation (SNSF) R’Equip grant 316030-139220, a SNSF Assistant Professorship grant PP00P3-144874, a Swiss Cancer League grant, the PhosphonetPPM SystemsX grant, and funding from the European Research Council (ERC) under the European Union’s Seventh Framework Programme (FP/2007-2013)/ERC Grant Agreement n. 336921. D Schapiro is supported by the Forschungskredit of the University of Zurich, grant FK-74419-01-01. HWJ and D Schulz are supported by European Molecular Biology Organization (EMBO) Long Term Fellowships co-funded by the European Commision (LTFCOFUND2013 and 2014), grants ALTF-711 2015 and ALTF-970 2014, respectively.

## Materials and Methods

### Preparation and staining of breast cancer tissue specimens

Formalin-fixed paraffin-embedded tissue samples from patients treated at the University Hospital Zurich between 1991 and 2005 were retrieved from the archives of the Institute of Surgical Pathology. This project was approved by the local Commission of Ethics (ref. no. StV 12-2005). H&E stained sections of all tumors were re-evaluated by a pathologist for their suitability for tissue microarray construction prior to array construction as previously described^22^.

Tissues were stained as previously described^10^. Briefly, tissue sections were de-waxed overnight in xylene and rehydrated in a graded series of alcohol (ethanol:deionized water 100:0, 90:10, 80:20, 70:30, 50:50, 0:100; 5 min each). Heat-induced epitope retrieval was conducted in a water bath at 95 °C in Tris-EDTA buffer at pH 9 for 20 min. After immediate cooling, the microarrays were blocked with 3% BSA in TBS for 1 hour. Samples were incubated overnight at 4 °C in primary antibody at 7.5 g/L diluted in TBS/0.1% Triton X-100/1% BSA (clones in Supplementary Table 1). Panel design and antibody database management was done in AirLab^23^. Samples were then washed twice with TBS/0.1% Triton X-100 and twice with TBS and dried before imaging mass cytometry measurement.

### Imaging mass cytometry

Antibody staining of tissue sections was quantified through the combination of laser ablation using a modified ArF excimer GeoLas C laser system (Coherent) to ablate tissues in a rastered pattern at 20 Hz and direct aerosol transportation of the sample to a CyTOF mass cytometer (Fluidigm) as described previously^24^. All raw data processing was performed using in-house Matlab routines as described and provided previously^13^.

### Segmentation

Segmentation was performed using Ilastik 1.1.9^25^ and CellProfiler 2.1.1^16^. Ilastik was used to classify pixels into three classes (nuclei, membrane, and background) and to generate probability maps. CellProfiler was used to segment probability maps to generate segmentation masks. A combination of channels was used to classify the background and membrane^26^. These masks were combined with the individual tiff files to extract single-cell information from each individual image.

### Single-cell feature extraction

histoCAT uses Matlab’s regionprops function to extract shape and pixel value measurements. Additionally, by step-wise pixel expansion, histoCAT creates a network of neighbors surrounding each cell at a range of defined distances. In most cases, expansion in the range of 1 to 6 pixels was chosen. Cells were expanded using a rectangular membrane shape. All cells within the defined range were considered neighbors. The distances between centroids were used to define cell-cell distances. The number of neighbors and the percent of membrane in contact with a neighboring cell were both determined with modified CellProfiler 1.0 modules^16^.

### Data transformation

Raw measurements were used for the presented data; however histoCAT offers arcsinh transformation with a variable cofactor input.

### Normalization

All images were segmented and single-cell measurements were extracted from all available channels using the mean pixel values for each segmented cell. The presented data were not normalized, but histoCAT features Z-score normalization across all samples or across a subgroup as a module. We used 99^th^ percentile normalized data for t-SNE and Phenograph as suggested^17,20^.

### histoCAT

histoCAT can be downloaded either as a Matlab 2014b app or as a stand-alone application for Mac, Windows, or Linux on the project page https://github.com/BodenmillerGroup/histoCAT. Documentation, user manual, and development versions of histoCAT can also be found on the project page. All modules, if not differently stated, were written in Matlab 2014b, and the GUI was designed in Matlab 2014b using Matlab’s GUI development environment (GUIDE). histoCAT is built modularly to enable addition of new features without need for changes to the existing structure. In general, features in histoCAT must include only two basic scripts: callback from the GUI and the script executing the function. The main functions are not linked to the GUI and can be run independently.

All data necessary to perform any function or stored for the current session can be retrieved from the GUI handles or included manually without the GUI. Throughout a session, the data are kept in the fcs-format structure. There is one main matrix containing a column for each channel and a row for each individual cell of each image. This matrix is continuously updated during the session and will therefore also contain the custom gates and channels. The corresponding channel names for each image are saved in a cell array. All individual tiff files and corresponding masks are stored in a multidimensional matrix structure.

### bh-t-SNE

We used the Barnes-Hut t-SNE implementation in histoCAT^17^. Data were 99^th^-percentile normalized before the analysis, and we used the default t-SNE parameters (initial dimensions: 110; perplexity: 30; and theta: 0.5). The random seeds for the individual runs can be recorded.

### PhenoGraph

PhenoGraph version 0.2 was used^15^. Data were 99^th^-percentile normalized before the analysis, and default parameters with nearest neighbors of 75 were used. This parameter was chosen based on prior knowledge of the underlying cell types. Lower values for nearest neighbors result in an over clustering and higher values an under clustering. The random seeds for the individual runs can be recorded.

### Neighborhood analysis – permutation test

Neighborhood analysis uses basic statistical methods to find significantly enriched interactions between or within cell phenotypes. First, cells are manually or automatically classified. Manual classification can be done by manual gating on biaxial/t-SNE plots. Automatic classification uses the PhenoGraph^15^ algorithm. PhenoGraph consistently performs well for datasets with multiple cell populations^14^.

Once classified, pairwise interactions at a distance of 6 pixels between and within cell phenotypes are calculated for each single cell with its neighbors. A neighbor is defined as a cell within the pixel distance selected during the loading process. Pairwise interactions between and within cell phenotypes are compared to a random distribution using a permutation test. This test provides us with a p-value for each one-tailed test. These p-values represent the likelihood of a neighborhood interaction being enriched or delimited in comparison to a random neighborhood. Equation 1 describes our approach using a permutation test with Monte Carlo sampling. We run this test twice to calculate the p-value for each tail.

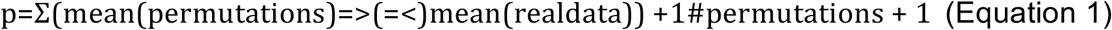

### Neighborhood analysis – Validation

The following simple examples demonstrate the validity of the neighborhood module in detecting cell neighborhoods deviating from randomness (Supplementary Figure 6). These artificial images were constructed in a simple chessboard pattern to visualize the validation. Complex structured synthetic datasets were validated but are not shown. The presented examples were used as a unit test in histoCAT to ensure the neighborhood analysis runs correctly on all platforms. The test dataset contained three different “phenotypic” clusters of cells, constructed by PhenoGraph: green represents cells of cluster 1, blue represents cells of cluster 2, and red cells are part of cluster 3.

All of the following examples were run at a pixel-expansion of 4 for 99 permutations and with a significance cut-off of 0.05 for the p-value. The hierarchically clustered heatmaps display the cell cluster interactions on the x-axis and the four test images on the y-axis. No interactions involving cluster 2 are observed. Those columns were automatically cut out of the visualization since their interaction frequencies did not significantly deviate from randomness. Thus the focus here is on clusters 1 and 3.

For image 1 the interaction between cells of cluster 3 with themselves (3→3) is displayed red in the corresponding row of the hierarchically clustered heatmap, indicating that this interaction occurs significantly more frequently in the actual test image than in the same image with randomly permuted cell labels (Supplementary Fig. 6a). This result is clearly supported by the bulk of cluster 3 cells in image 1. Random permutations of the cell labels are likely to distribute cells of cluster 3 among those of cluster 1. This also leads to a significantly fewer cluster 1 cells neighboring cluster 3 cells in the permuted images compared to our real image, hence the blue 1→3 interaction in the hierarchically clustered heatmap (Supplementary Fig. 6a).

The alternating pattern of cells of clusters 1 and 3 in image 2 prevents cells of the same cluster from being in each other’s neighborhood (Supplementary Fig. 6b). Therefore the interactions 1↔1 and 3↔3 are significantly less frequent than in random permutations of the cluster labels.

Image 3 reaches the significance cut-off for none of the cell-type neighbor interactions, despite the fact that two of the cells of cluster three are neighbors (Supplementary Fig. 6c). This is expected and visualizes the effect of the cut-off.

The hierarchically clustered heatmap of image 4 shows that the interactions between cells of cluster 1 and cells of cluster 3 occur significantly more often (in both directions) than expected from a randomly shuffled image, as is clearly visible in the test image on the left where cells of cluster 1 always neighbor a cell of cluster 3 and vice versa (Supplementary Fig. 6d).

## Contributions

D Schapiro, HWJ, and BB conceived of the project and software. HWJ, CG, and RCF collected samples and validated antibodies. ZV assembled, classified, and provided tumor samples. HWJ completed the staining and image acquisition. D Schapiro, SR, and JF wrote the code. D Schapiro, HWJ, and D Schulz tested software on multiple data sources. D Schapiro, HWJ, and VZ analyzed the images and single cell data. D Schapiro, HWJ, and BB prepared the figures and wrote the manuscript. BB directed the project.

